# Comparing Nonlinear Trajectories Across Brain Networks: A Key to Understanding Complex Brain Dynamics

**DOI:** 10.64898/2026.07.24.740420

**Authors:** Masoud Seraji, Sarah Shultz, Qiang Li, Zening Fu, Vince D Calhoun

## Abstract

This study explores the nonlinear developmental trajectories of brain networks in neurotypically developing infants during their first six months. Using a longitudinal dataset of 137 resting-state functional MRI scans from 74 infants, we analyzed five spatial metrics across 13 intrinsic connectivity networks, including motor, visual, subcortical, and prefrontal networks. A cubic model was specifically employed to capture distinct linear and non-linear trends in the networks’ developmental patterns, allowing for the identification of significant differences in cubic, quadratic, and linear slope parameters across networks. This model choice was driven by the need to examine how specific non-linear components (e.g., inflection points and acceleration rates) uniquely characterize each network’s trajectory, which a generalized approach might smooth out without pinpointing such network-specific features. Notably, the subcortical network exhibited a distinct cubic growth pattern, while secondary motor and visual networks showed pronounced quadratic variations, suggesting network-specific shifts in spatial organization and connectivity. These findings highlight the unique maturation timelines and interactions between functional systems, such as early sensory-motor coordination and later cognitive integration. The results underscore the importance of network-specific growth patterns, providing deeper insights into how infant brain networks evolve and interact to support emerging cognitive and behavioral functions.

## 1. Introduction

Nonlinear changes in brain structure or function (e.g., changes with age during development) are commonly studied, revealing complex changes in brain activity over time (Cao et al., 2017b; Faghiri et al., 2019; Gao et al., 2015a; Khalilian et al., 2024; Monroe et al., 2022; Sørensen et al., 2021) but little attention has been given to comparing these trajectories across networks. This gap is significant because brain networks do not function in isolation; they work in concert to support cognition, perception, and behavior (Pessoa, 2023). The core idea of comparing nonlinear trajectories across brain networks is to understand how different parts of the brain develop at varying rates and interact as a system. This can be observed by comparing the varying shapes of their nonlinear trajectories. In this study, we applied a cubic model to characterize these trajectories, offering an effective compromise between model interpretability and analytical flexibility. In a cubic model, each term contributes distinct nonlinear behavior. The cubic term (*a*_1_*x*^3^) introduces an S-shaped curve, allowing for more complex dynamics like inflection points, where the direction of curvature changes. This term introduces asymmetry in the curve and can represent multiple turning points, with stronger effects at the extremes as the value of a increases. This enables us to model complex, non-linear developmental patterns—such as sharp directional shifts or multi-phase trajectories (e.g., an early increase followed by a steep decline, or a plateau preceding a decrease)—that simpler models fail to capture. The quadratic term (*a*_2_ *x*^2^) adds parabolic curvature, allowing the relationship to deviate from a straight line and capture acceleration or deceleration (its direction depends on the sign and range of *a*_2_ ). This term produces symmetric curves, effectively modeling systems with peaks or valleys, where the rate of change slows before reversing, such as in growth or decay processes. Finally, the linear term (*a*_3_ *x*) provides a straight-line relationship between input and output, contributing no nonlinearity by itself but interacting with the cubic and quadratic terms (Bostock and Chandler, 2002; Hughes-Hallett, 2018).

Our view is that different coefficients associated with each term in a cubic model (*a*_1_, *a*_2_, and *a*_3_) play an important role in the trajectory, reflecting the distinct rates of change in each network. By examining and comparing these coefficients across different networks, we can assess whether systems exhibit similar nonlinear behaviors or if unique variations emerge. This type of comparison helps uncover whether the networks mirror each other’s dynamics or diverge in their responses to changes, indicating periods of rapid or slow development (Elhakeem et al., 2022). In addition, analyzing nonlinear patterns across networks provides deeper insights into global brain dynamics. For example, the development of the default mode network (DMN) and the frontoparietal network (FPN) can be modeled using the terms *a*_1_, *a*_2_, and *a*_3_ of a cubic trajectory. The DMN, which reaches an adult □like activation profile earlier in development (Chen et al., 2023), is therefore expected to display a stronger linear term that reflects relatively stable, incremental growth across infancy and early childhood. In contrast, the FPN - whose maturation lags behind (Chen et al., 2023)-could exhibit pronounced quadratic or cubic coefficients, capturing a later phase of accelerated expansion and rising configurational complexity. This developmental offset is consistent with evidence that a mature DMN provides a scaffold for emerging executive systems: Chen □et□al.□(2023) reported that posterior □ cingulate DMN nodes attain adult□ level patterns before dorsolateral prefrontal FPN nodes, and that coupling between these two networks mediates gains in working□memory performance (Chen et al., 2023).

To the best of our knowledge, this is the first study to systematically compare nonlinear trajectory parameters across multiple brain networks in infants, a method we term cross-network nonlinear trajectory characterization. In this initial work, we employed a relatively simple cubic model to explore age-related changes across five distinct metrics within 13 brain networks. By fitting a cubic model to each network and then statistically contrasting the linear, quadratic, and cubic coefficients between every pair of networks, we captured how their growth curves differ in magnitude and shape. This *cross-network* comparison goes beyond within-network characterization and reveals which systems mature earlier, later, or more abruptly than others. These variations revealed meaningful insights into how different brain networks evolve over time, highlighting distinct growth patterns. This novel approach offers a fresh perspective on comparing developmental patterns across networks, highlighting differences in their maturation timelines and suggesting potential implications for how systems may align during development. While our framework does not directly measure inter-network coupling, contrasting nonlinear trajectories provides indirect insight into which networks may mature earlier and potentially scaffold later-developing systems.

## 2. Materials and Methods

### 2.1 Imaging dataset

This study collected longitudinal resting-state functional magnetic resonance imaging (rs-fMRI) data from 74 neurotypically developing infants (43 males, 31 females) aged 0 to 6 months. A total of 137 scans were obtained, with infants undergoing up to three scans each between birth and 6 months, as illustrated in Fig. 1. Infants were typically developing, with no family history of autism in up to 3^rd^ degree relatives, no family history of developmental delay in 1^st^ degree relatives, and no perinatal complications, neurological □/□sensory or genetic conditions, seizures, or MRI contraindications. The analysis used corrected age—calculated by adjusting for deviations from the standard 40-week gestational period—rather than chronological age (Glass et al., 2015). Informed consent was obtained from all legal guardians, and the study received ethical approval from Emory University. The details of the data collection procedures are described in detail in our previous study (Seraji et al., 2025a).

**Fig. 1.**
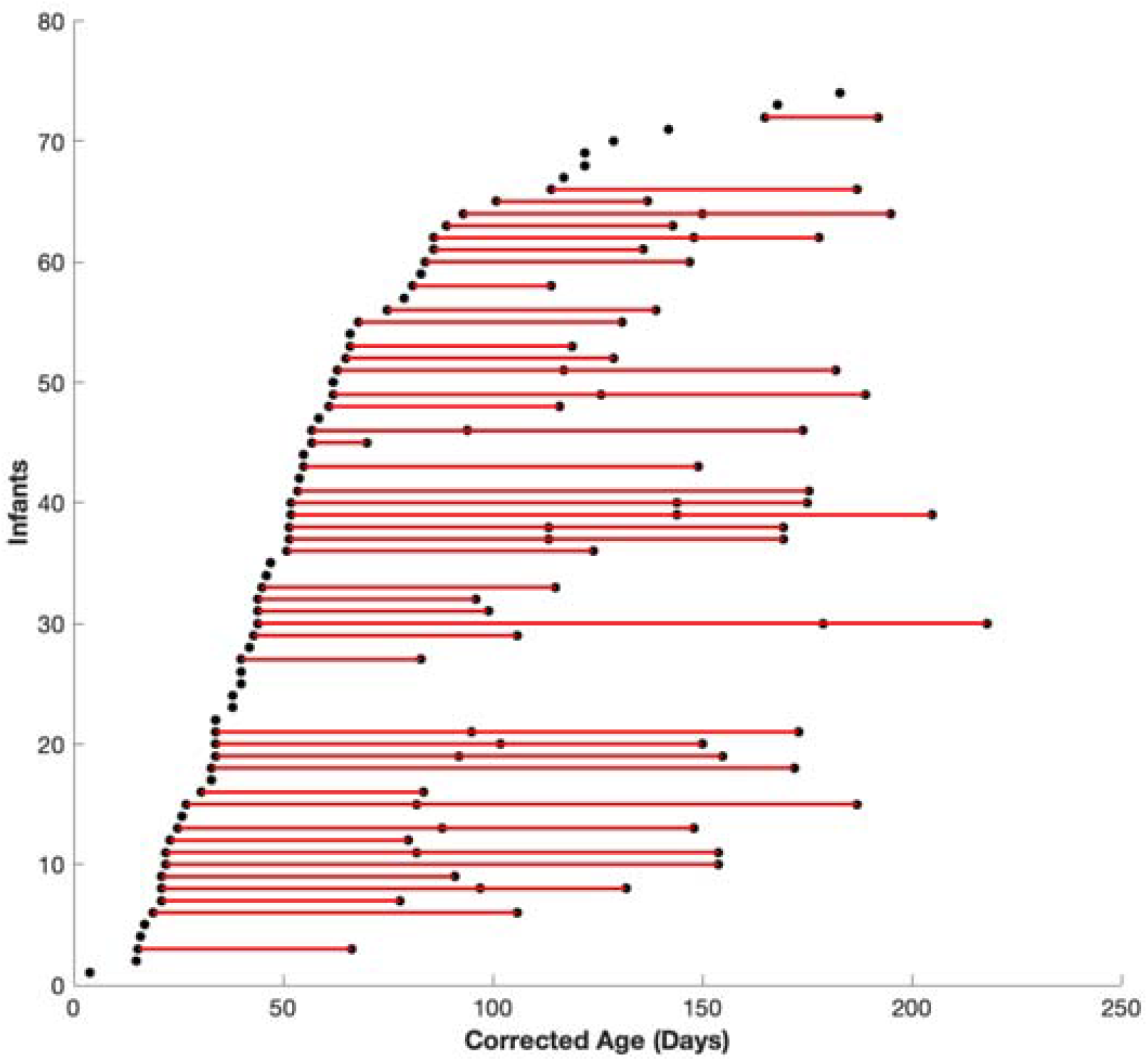
The distribution of scans by corrected age for all participants. Each dot represents a single scan from a participant, with dots connected by lines indicating all longitudinal scans obtained from the same individual. The data were collected using a non-uniform longitudinal sampling design, with each infant undergoing up to three scans at pseudorandom time points between birth and 6 months. In total, 137 scans were acquired, with an average interval of 1.4 days between scans (SD = 1.8).

### 2.2 Preprocessing

The preprocessing of the fMRI data involved several steps. First, the initial 16 volumes were discarded to achieve magnetization equilibrium. Head motion correction was performed using the mcflirt function in FSL, aligning all volumes to the first one. The single-band data acquired with phase encoding in AP and PA direction to estimate the susceptibility-induced off-resonance field, which was used to correct the distortion in the multi-band resting-state fMRI data, followed by a slice-timing correction. A two-step spatial normalization process was applied: initially, the infant fMRI data were co-registered and normalized to a 3-month T1 template from the Baby Connectome Project (BCP) (Chen et al., 2022) to capture developmental dynamics, and then further aligned to the Montreal Neurological Institute (MNI) space using an EPI template (Fu et al., 2023). Finally, spatial smoothing was applied using a 6 mm Gaussian kernel.

### 2.3 Data Analysis

Quality control involved excluding scans with poor alignment to MNI space by comparing individual masks with the group mask (2 scans were excluded; See (Seraji et al., 2025a) for more details). Group independent component analysis (ICA) was conducted using the GIFT toolbox to identify large-scale brain networks (Calhoun et al., 2001; Iraji et al., 2021). The process began with subject-specific principal component analysis (PCA) to reduce noise, enhance computational efficiency, and ensure each subject’s contribution was comparable. Thirty principal components (PCs) capturing the most variance were retained from each subject, with subject-level PCA emphasizing individual differences. These PCs were then concatenated, followed by group-level PCA to identify shared patterns across subjects. The top 20 group-level PCs were used as input for group ICA, following recommended practices for capturing large-scale networks (Iraji et al., 2016). The ICA was repeated 100 times with the Infomax algorithm to ensure stability through bootstrapping, and the most reliable components, identified by alignment with stable cluster centroids, were selected for further analysis. Finally, subject-specific large-scale brain networks were estimated using the group information guided (GIG)-ICA approach, ensuring individualized network mapping (Du et al., 2016). Further details of this analysis can be found in our previous paper (Seraji et al., 2025a).

### 2.4 Cross-network Non-linear Trajectory Characterization

After the ICA decomposition, each independent component was inspected for its spatial activation pattern; components whose patterns matched known functional motifs were labeled accordingly, yielding 13 large-scale brain networks: primary and secondary visual networks, subcortical network, cerebellum network, primary and secondary motor networks, attention network, default mode network, temporal network, auditory network, frontal-medial prefrontal (mPFC) network, frontal-dorsolateral prefrontal (dlPFC) network, and frontal-ventrolateral prefrontal (vlPFC) network.

We utilized several metrics to examine developmental changes in the spatial organization of brain networks, all derived from voxel intensities in spatial maps, which reflect each voxel’s contribution to the network. Network-averaged spatial similarity (NASS) quantifies the alignment between an individual’s spatial map and the group-level network map, with higher NASS values indicating closer conformity to shared spatial patterns and lower values suggesting more individual-specific variations. Network engagement range (NER) measures the variability in voxel contributions by calculating the difference between the highest and lowest voxel intensities, reflecting the range of engagement across the network. A broader range indicates greater variability, while a narrower range suggests a more uniform voxel contribution. Network strength captures the average intensity of all voxels within a network, indicating the level of voxel contribution; higher values reflect increased participation. This metric is computed by averaging the intensities of voxels exceeding a Z-score threshold of 1.96 (p = 0.05). Network size assesses the spatial extent of a network by counting the number of voxels that meet the same threshold, reflecting changes in the network’s spatial footprint over time. Network center of mass (NCM) evaluates the spatial configuration of a network by determining the center of mass

(COM) of activation clusters and calculating the weighted average distance of each voxel from the COM, providing insights into the network’s spatial distribution (Seraji et al., 2025a, 2025b).To examine developmental changes across networks, we assessed whether each metric exhibited linear and nonlinear age-related trends by fitting a cubic model across age for each of the five metrics within all 13 networks. The cubic model (*a*_1_*x*^3^ +*a*_2_ *x*^2^ +*a*_3_ *x* + *a*_4_) allowed us to capture both linear and nonlinear trends in developmental trajectories. For each metric and network, we extracted the cubic (*a*_1_ ), quadratic (*a*_2_ ), and linear slope (*a*_3_ ) parameters from the model. We modeled each network’s trajectory with a cubic polynomial using the same, shared design matrix (demeaned age terms). For every pair of networks, we compared the corresponding coefficients by taking their difference and computing a pooled standard error from the ordinary least squares fit. This standard error also accounts for cross-network residual covariance, so contrasts reflect shared noise as well as within-network variability. Each contrast was tested with a two-sided t-test (using the model’s residual degrees of freedom), and the resulting p-values were adjusted for multiple comparisons using false discovery rate (FDR) correction to ensure robust statistical significance. This comprehensive approach enabled us to characterize network-specific developmental trends and to contrast the relative magnitudes and directions of linear, quadratic, and cubic terms across networks, providing deeper insight into how the spatial organization of these 13 networks evolve during infancy.

## 3. Results

This section is divided into two key parts. In the first part, we applied a cubic model to fit our five selected metrics across all 13 networks, examining how each network behaves under this model. In the second part, we conducted a comparative analysis between the cubic, quadratic, and slope parameters across the networks, focusing on differences in their spatial metrics.

### 3.1 Non-Linear Developmental Trajectories in Brain Networks

To examine the non-linear trajectories for each metric within each network, we applied a cubic model *a*_1_*x*^3^ +*a*_2_*x*^2^ + *a*_3_*x* + *a*_4_ across age. Fig. 2 displays the fitted cubic curves for 13 networks: primary visual, secondary visual, cerebellum, primary motor, secondary motor, attention, default mode, temporal, auditory network, frontal-mPFC, frontal-dlPFC, and frontal-vlPFC.

**Fig. 2.**
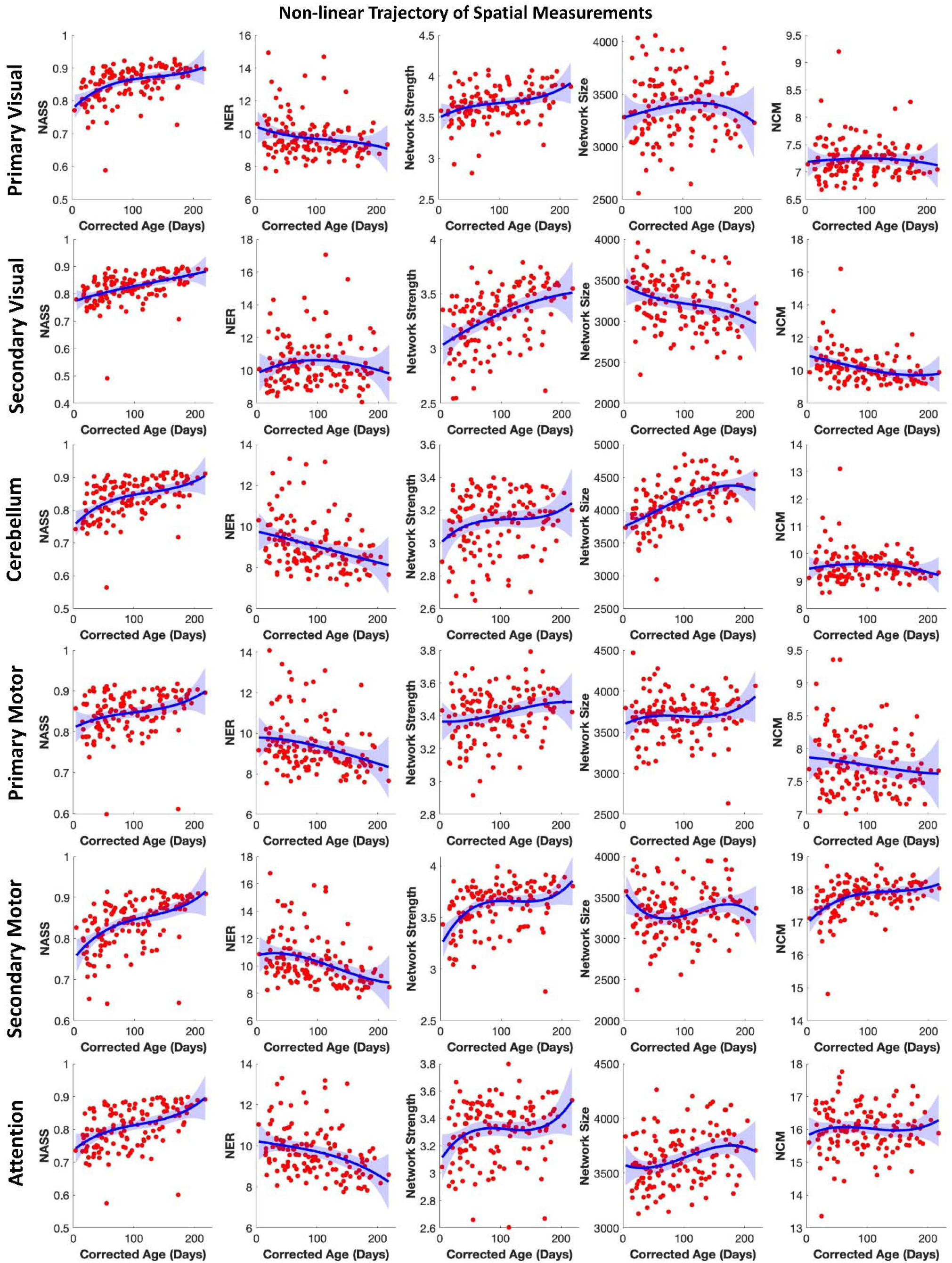
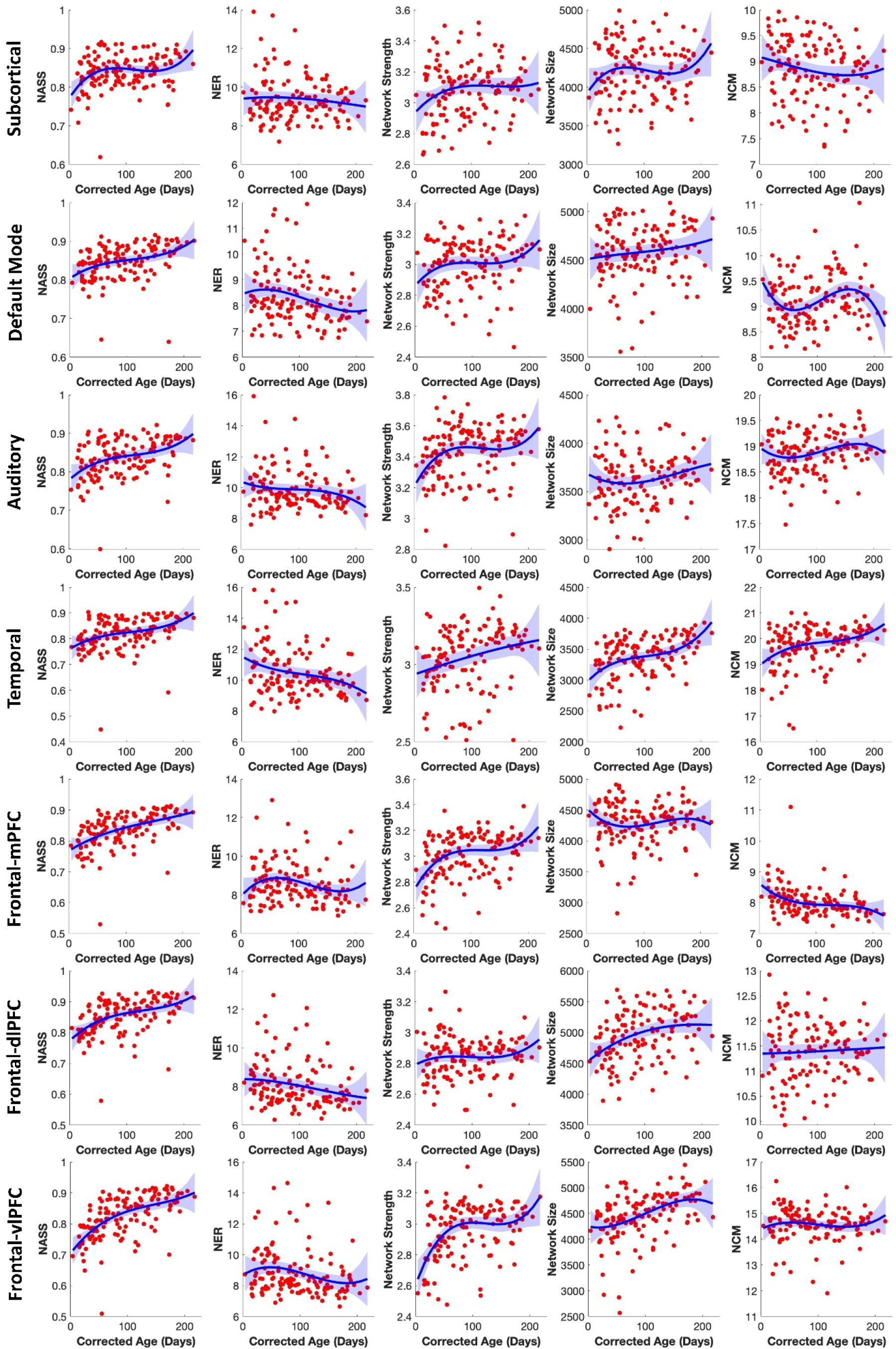
Non-linear trajectories of spatial measurements across age for various brain networks. Each row represents a different brain network, including the primary visual, secondary visual, cerebellum, primary motor, secondary motor, attention, subcortical, default mode, auditory, temporal, frontal-mPFC, frontal-dlPFC, and frontal-vlPFC networks. NASS: network-averaged spatial similarity; NER: network engagement range; NCM: network center of mass. Red dots correspond to individual scans for each subject, while the blue lines depict the group-averaged non-linear trajectories fitted to the data. Shaded areas around the fitted lines show 95% confidence intervals.

### 3.2 Comparative Analysis of Cubic, Quadratic, and Slope Parameters Across Networks

After fitting the cubic model and conducting the comparative analysis of the cubic, quadratic, and slope parameters across all networks, we evaluated the p-values associated with these differences (Δ’s) to determine statistical significance (See method). For enhanced clarity, Fig. 3 presents the results by visualizing −*log*_10_ (*p*) * *sign*(Δ) rather than raw p-values. This transformation highlights both the magnitude and direction of the differences more effectively, offering a clearer interpretation of significant variations between networks.

**Fig. 3.**
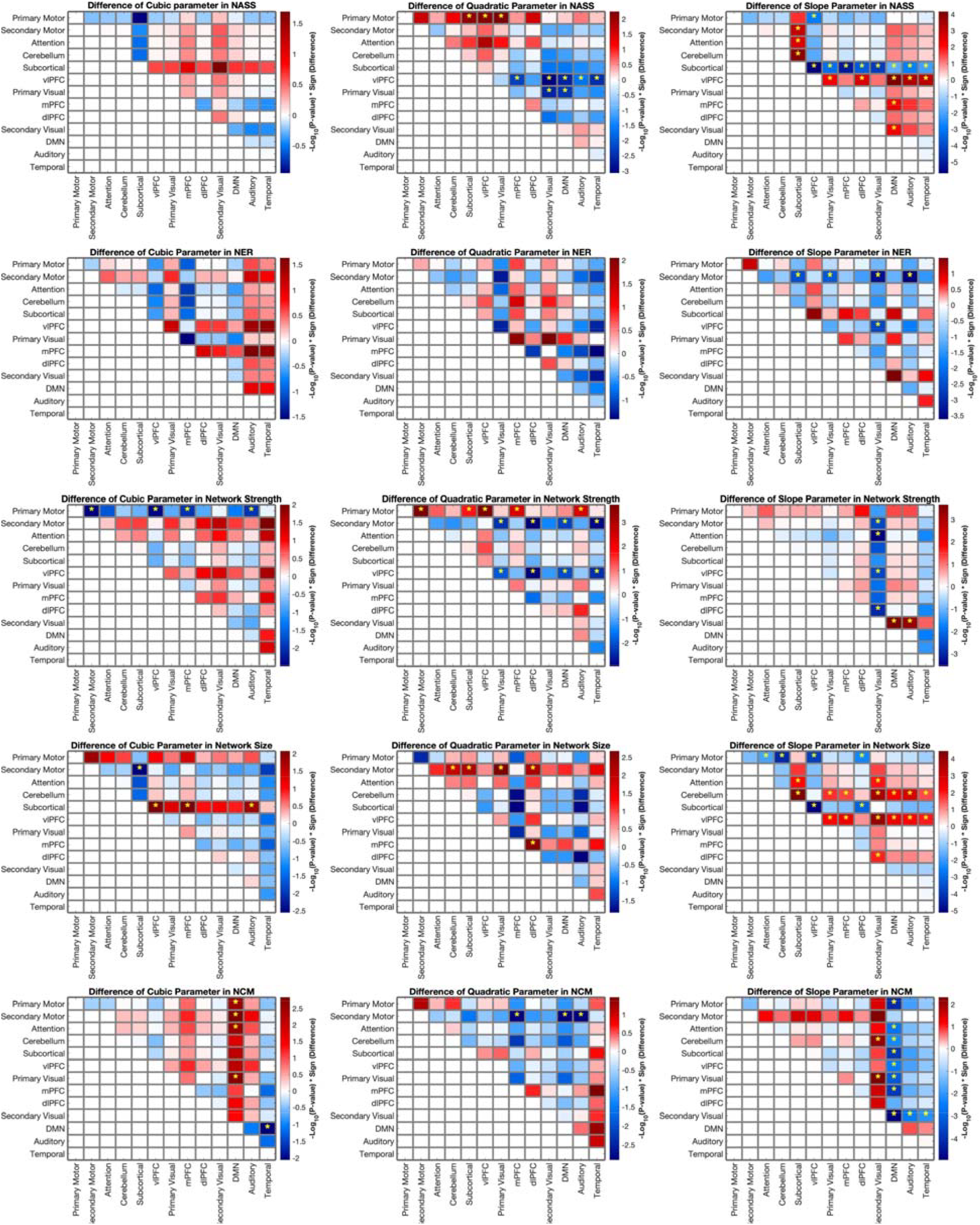
Comparison of cubic model coefficients across brain networks. Heatmaps display the “-log(p-value) ×sign(difference)” for the differences in cubic, quadratic, and slope parameters among all the identified brain networks. Each row represents a network, and each column represents the comparison network. Significant differences are highlighted with a yellow star, providing a clear visualization of how each network’s developmental trajectory differs from others in terms of nonlinear trends. The statistical analysis results were corrected for multiple comparisons using a 5% FDR.

As illustrated in Fig. 3, there are no significant changes in the cubic parameter of NASS between different networks. However, the quadratic and slope parameters showed clear cross-network differences. For quadratic, the primary motor and visual had significant difference with other networks (|t| > 2.31, p < 0.03). For the slope parameter that significant difference was between the subcortical network and all the other networks excep primary motor (|t| > 3.10, p < 0.02). Additionally, the vlPFC slope parameter significantly differs from half of the other networks. For NER, there are no significant differences in the cubic and quadratic parameters. Only a few slope contrasts were significant, primarily involving the secondary motor network compared with the primary visual, secondary visual, auditory, and subcortical networks (|t| > 3.09, p < 0.02).

For the network strength metric, pairwise contrasts revealed a small number of significant cubic effects, primarily involving the primary motor network, which differed from several other systems (e.g., secondary motor, vlPFC, mPFC, Auditory; |t| > 2.60, p < 0.03). In contrast, the quadratic term showed a broader pattern of cross-network differences: primary motor differed from multiple networks including secondary motor, subcortical, vlPFC, mPFC, and auditory, and the vlPFC network differed from dlPFC, primary visual, DMN, and temporal (|t| > 2.35, p < 0.04). The slope parameter was most distinctive for the secondary visual network, which showed significant differences from a subset of other systems—including secondary motor, vlPFC, DMN, and auditory (|t| > 3.03, p < 0.03).

The cubic parameter of network size shows significant differences, primarily between the subcortical network and other networks (|t| > 2.44, p < 0.04). The quadratic (|t| > 2.53, p < 0.03) and slope (|t| > 2.74, p < 0.04) parameters for this metric also exhibit significant differences between several networks, as shown in Fig. 3. For the NCM, the cubic parameter indicates significant differences mainly between the default mode network and other networks (|t| > 3.01, p < 0.03), including the primary and secondary motor, attention, primary visual, and temporal networks. The quadratic (|t| > 3.26, p < 0.005) and slope (|t| > 2.81, p < 0.04) parameters of the NCM show differences primarily between the secondary motor, secondary visual, default mode networks, and other networks, with detailed differences illustrated in Fig. 3. Overall, the figure illustrates how different brain networks exhibit unique non-linear developmental trajectories, with some networks showing distinct differences compared to others and some showing more similarities.

## 4. Discussion

Our findings highlight the non-linear developmental trajectories of brain networks during the first six months of life, emphasizing that brain networks exhibit distinct, network-specific maturation patterns. These results align with previous studies that underscore how different brain networks mature at varying rates and through diverse patterns of connectivity changes (Cao et al., 2017b; Gao et al., 2015a) .

Significant differences in slope parameters indicate that infant functional networks differ in the rate at which their spatial organization becomes refined across age. Networks with steeper positive slopes show greater age-related increases in NASS, meaning that their spatial maps become more aligned with the group-level network pattern more rapidly. In our trajectories, subcortical and vlPFC regions show more sharply rising curves, suggesting accelerated early spatial refinement, whereas other networks show flatter trajectories, indicating slower or more gradual maturation. Together, these cross-network differences suggest that early brain network maturation is heterochronous, with different systems following distinct developmental tempos. While these results do not directly identify the biological mechanisms driving these differences, they demonstrate variation in the timing and rate of spatial maturation, consistent with prior evidence for heterogeneous and nonlinear developmental trajectories across brain systems (Cao et al., 2017b; Gao et al., 2015a). Across networks, network strength increased with age, but the trajectories differed in both rate and curvature, revealing meaningful differences in maturation timing. Most of the networks exhibit an early, rapid increase during the first ∼80–100 days, followed by a more gradual rise and in some cases a late acceleration approaching ∼180–200 days. The significant differences in the quadratic parameter refine this interpretation: networks such as the secondary motor and vlPFC display stronger curvature, suggesting more pronounced early acceleration and earlier inflection points, whereas primary motor and primary visual networks follow a more gradual, linear-like increase. The slope parameter highlights networks with faster overall strengthening—most notably the secondary visual network, which shows a steeper trajectory than nearly all others, indicating a more rapid and sustained increase across infancy.

The subcortical network shows a steep early increase in network size over the first ∼60–80 days, followed by a slowing/near-stable phase from roughly 80–150 days. After that, the curve shows a modest re-acceleration between ∼150–200 days, indicating renewed but more gradual growth rather than a true plateau.The unique cubic parameter for network size in the subcortical network suggests distinct phases of rapid growth and reorganization compared to other networks (Alex et al., 2024). Subcortical regions, which play foundational roles in early development, may experience accelerated structural changes before stabilizing (Grotheer et al., 2022). In addition, steeper slopes in cerebellum and vlPFC networks reflect a rapid early expansion in size during the first ∼2–4 months, followed by a clear flattening, suggesting that much of their growth occurs in the middle of this six months. In contrast, quadratic effects were sparse and localized, indicating that most networks followed broadly similar curvature while only a few showed distinct tempo of expansion. The secondary motor network exhibited the strongest quadratic effect because its size trajectory showed a distinctive U-shaped pattern—an early decline followed by later re-expansion—whereas most other networks changed more monotonically across the 0–6 month window. Across networks, NCM showed relatively modest nonlinear change, but two networks displayed distinct developmental signatures captured by the model coefficients. The DMN was the only network with significant cubic differences from several others, reflecting its uniquely tri-phasic trajectory: an early decline in NCM during the first month, followed by a mid-infancy rebound around 40–140 days, and a late downturn toward 200 days. This S-shaped pattern requires a cubic term and also explains why the DMN shows significant slope differences relative to both increasing and decreasing networks—its net slope is near zero because opposing phases counterbalance each other. In contrast, quadratic differences were concentrated in the secondary motor network. Together, these findings highlight that NCM captures network-specific developmental motifs—with the DMN showing multi-phase restructuring rather than a uniform maturation pattern across the brain.

In summary, our results underscore that early brain network development is highly heterogeneous and non-linear, with each network following its own maturation timetable (Cao et al., 2017a; Wang et al., 2023). This aligns with established developmental principles: primary sensory and motor circuits tend to mature earlier and more rapidly, whereas higher-order associative networks (e.g. the default mode network) evolve more gradually (Gao et al., 2015b). For instance, newborn sensorimotor networks are comparatively adult-like and strengthen quickly, while networks subserving complex cognitive functions remain immature at birth and only synchronize over time (Gao et al., 2015b). We observed early surges in the organization of subcortical, cerebellar, and ventrolateral prefrontal networks—critical systems for basic sensorimotor and regulatory functions—consistent with the explosive structural growth and connectivity in these regions during infancy (the cerebellum alone doubles in volume by ∼3 months) (Holland et al., 2014). Such rapid early gains likely reflect biological processes like initial circuit formation and myelination, which are known to progress unevenly across brain regions in the first half-year (Grotheer et al., 2022). By contrast, the default mode network displayed a unique multi-phase trajectory, suggesting that some higher-order networks undergo complex reorganization rather than a steady linear increase. This pattern is reminiscent of broader neurodevelopmental phenomena in which periods of synaptic overgrowth are followed by pruning and refinement (Petanjek et al., 2011). Taken together, these findings provide a nuanced picture of infant brain maturation, highlighting that developmental changes occur at different rates and in different forms across networks. Appreciating these network-specific trajectories is crucial for understanding typical brain development and may inform early identification of atypical patterns, especially since the first months of life represent a time of both high neuroplasticity and vulnerability to developmental disorders.

## 5. Conclusion

In the first six postnatal months, brain networks follow distinct, non-linear developmental trajectories. Subcortical and cerebellar systems exhibit rapid early expansion, while higher-order networks like the DMN and vlPFC mature more gradually and in multi-phase patterns. Significant differences in slope, quadratic, and cubic terms reveal diverse growth tempos—such as the U-shaped expansion of the secondary motor network and the tri-phasic restructuring of the DMN. These findings underscore the asynchronous nature of early brain development, where early-maturing systems may scaffold later-emerging functions. This normative characterization provides a foundation for identifying atypical trajectories in future work.

